# Specific plasticity loci and their synergism mediate operant conditioning

**DOI:** 10.1101/2021.12.02.470828

**Authors:** Yuto Momohara, Curtis L. Neveu, Hsin-Mei Chen, Douglas A. Baxter, John H. Byrne

## Abstract

Despite numerous studies examining the mechanisms of operant conditioning (OC), the diversity of plasticity loci and their synergism have not been examined sufficiently. In the well-characterized feeding neural circuit of *Aplysia*, appetitive OC increases neuronal excitability and electrical coupling among several neurons. Here we found OC decreased the intrinsic excitability of B4 and the strength of its inhibitory connection to a key decision-making neuron, B51. The OC-induced changes were specific without affecting the B4-to-B8 inhibitory connection or excitability of another neuron critical for feeding behavior, B8. A conductance-based circuit model indicated certain sites of plasticity mediated the OC phenotype more effectively and that plasticity loci acted synergistically. This synergy was specific in that only certain combinations of loci synergistically enhanced feeding. Taken together, these results suggest modifications of diverse loci work synergistically to mediate OC.

**Significance Statement:** The diversity and synergism of plasticity loci mediating operant conditioning (OC) is poorly understood. Here we found that OC decreased the intrinsic excitability of a critical neuron mediating *Aplysia* feeding behavior and specifically reduced the strength of one of its inhibitory connections to a key decision-making neuron. A conductance-based computational model indicated that the known plasticity loci showed a surprising level of synergism to mediate the behavioral changes associated with OC. These results highlight the importance of understanding the diversity, specificity and synergy among different types of plasticity that encode memory. Also, because OC in *Aplysia* is mediated by dopamine (DA), the present study provides insights into specific and synergistic mechanisms of DA-mediated reinforcement of behaviors.

## Introduction

Appetitive operant conditioning (OC) increases expression of specific behaviors due to the association of the behavior with rewards (Skinner, 1981; Thorndike, 1933). This form of learning is ubiquitous (for reviews see Martin-Soelch et al., 2007; Nargeot and Puygrenier, 2019) and is involved in addiction (Koob and Volkow, 2010). The neuronal mechanisms of appetitive OC include changes in synaptic strength and intrinsic excitability of individual neurons (for review see Mozzachiodi and Byrne, 2010; Cox and Witten, 2019; Nargeot and Puyrenier 2019). Learning likely involves diverse plasticity mechanism that work synergistically (for example, see Gao et al., 2012). However, the diversity of plasticity loci, and how they might act synergistically to mediate OC remains relatively unexplored.

To address this issue, we exploited the technical advantages of *Aplysia* feeding behavior, which is mediated by a central pattern generator (CPG) located primarily in the buccal ganglia (Church and Lloyd, 1994; Elliott and Susswein, 2002; Cropper et al., 2004) and can be modified by appetitive OC (Brembs et al., 2004; Baxter and Byrne, 2006; Nargeot and Simmers, 2011). The CPG generates at least two mutually exclusive behaviors, egestion and ingestion. During OC, ingestion behavior is reinforced by contingent presentation of food (*in vivo*) or stimulation of dopaminergic afferents (*in vitro*) (Nargeot et al., 1997, 2007; Brembs et al., 2002). Previously identified neuronal correlates of OC include increases in the excitability and electrical coupling of neurons that initiate the behavior (Nargeot et al., 2009; Sieling et al., 2014; Costa et al., 2020), and increased excitability of a plateau generating neuron, B51, involved in selection of ingestion (Nargeot et al., 1999a,b; Lorenzetti et al., 2006, 2008; Mozzachiodi et al., 2008).

Surprisingly, to date all of the correlations of OC in the *Aplysia* feeding system are associated with either increases in intrinsic neuronal excitability or electrical coupling. These results raise several important questions. First, does OC decrease excitability of neurons? Second, does OC modify chemical synaptic transmission? Third, are modifications of some neurons or synapses more effective than others? Fourth, in what ways would such a diverse array of neuronal plasticity act synergistically to alter behavior? To address these issues, we examined whether OC produces complementary modifications in both the excitability and strength of inhibitory connections of B4, a decision-making neuron that suppresses ingestion, in part, by its inhibitory synaptic connections to B8 and B51 (Fig. 2*A*) (Plummer and Kirk, 1990; Kabotyanski et al., 1998; Sasaki et al., 2009; Dacks et al., 2012). We then investigated the behavioral consequences of any observed modifications by including these changes in a conductance-based model of the CPG and examining whether they synergistically enhanced feeding.

## Materials and Methods

Three distinct but complementary approaches were used in this study: 1) an *in vitro* analogue of operant conditioning in isolated preparations of buccal ganglia dissected from naïve animals, 2) a single-cell analogue of OC in cell culture with neurons isolated from buccal ganglia of naïve animals; and 3) a conductance-based model of the CPG to assess the relative contributions of the changes in excitability and synaptic strength to the generation of BMPs.

### Animals

*Aplysia californica* (30-70 g) were obtained from the National Resource for *Aplysia* (University of Miami, FL), Alacrity Marine Biological Specimens (Redondo Beach, CA), and Marinus (Westchester, CA). Animals were housed individually in perforated plastic cages in aerated seawater tanks at a temperature of 15 °C, and were fed ∼1 g of dried seaweed three times per week.

### Classification of buccal motor patterns (BMPs)

Feeding behavior can be broadly classified into behaviors that lead to the ingestion or rejection of food. Both ingestion and rejection consist of two phases that involve an outward (protraction) and inward movement (retraction) of the radula, a grooved tongue-like structure. Radula closure during the retraction phase is a distinguishing feature of ingestion behavior. Rhythmic movements of the radula during feeding are controlled by a CPG within the buccal ganglia, which continues to produce patterned motor activity in isolated ganglia. This patterned motor activity (i.e., buccal motor patterns, BMPs) was monitored *in vitro* by extracellular recordings of nerves I2 n, R n.1 and n.2,1 (Fig. 1*A,* 1*B2*). Similar to previous studies (Nargeot et al., 1997), fictive protraction and retraction was designated manually based on activity in I2 n. and n.2,1 respectively. During post hoc analysis, radular closure was estimated automatically by a custom-built spike detection algorithm analyzing recordings of R n.1 (Fig. 1*A*) or B8 in preparations that included intracellular recordings from B8 (Fig. 2*B*). The spike detection consisted of a simple threshold that was manually set for each nerve recording, to be above the smallest R n.1 spike during protraction that was greater than spikes (likely noise) outside of BMP activity. Easily discernable large unit R n.1 spikes occurring outside of BMP activity were ignored when setting the threshold. Patterns with >50% of the total duration of R n.1 activity occurring during retraction were classified as iBMPs (Nargeot et al., 1997, 1999a,b,c; Brembs et al., 2002; Mozzachiodi et al., 2008) and for simplicity, all other patterns were classified as rejection buccal motor patterns (rBMPs). For each preparation, BMP classification was carried out by an individual blind to the stimulus protocol used (OC vs. control).

**Figure 1.**
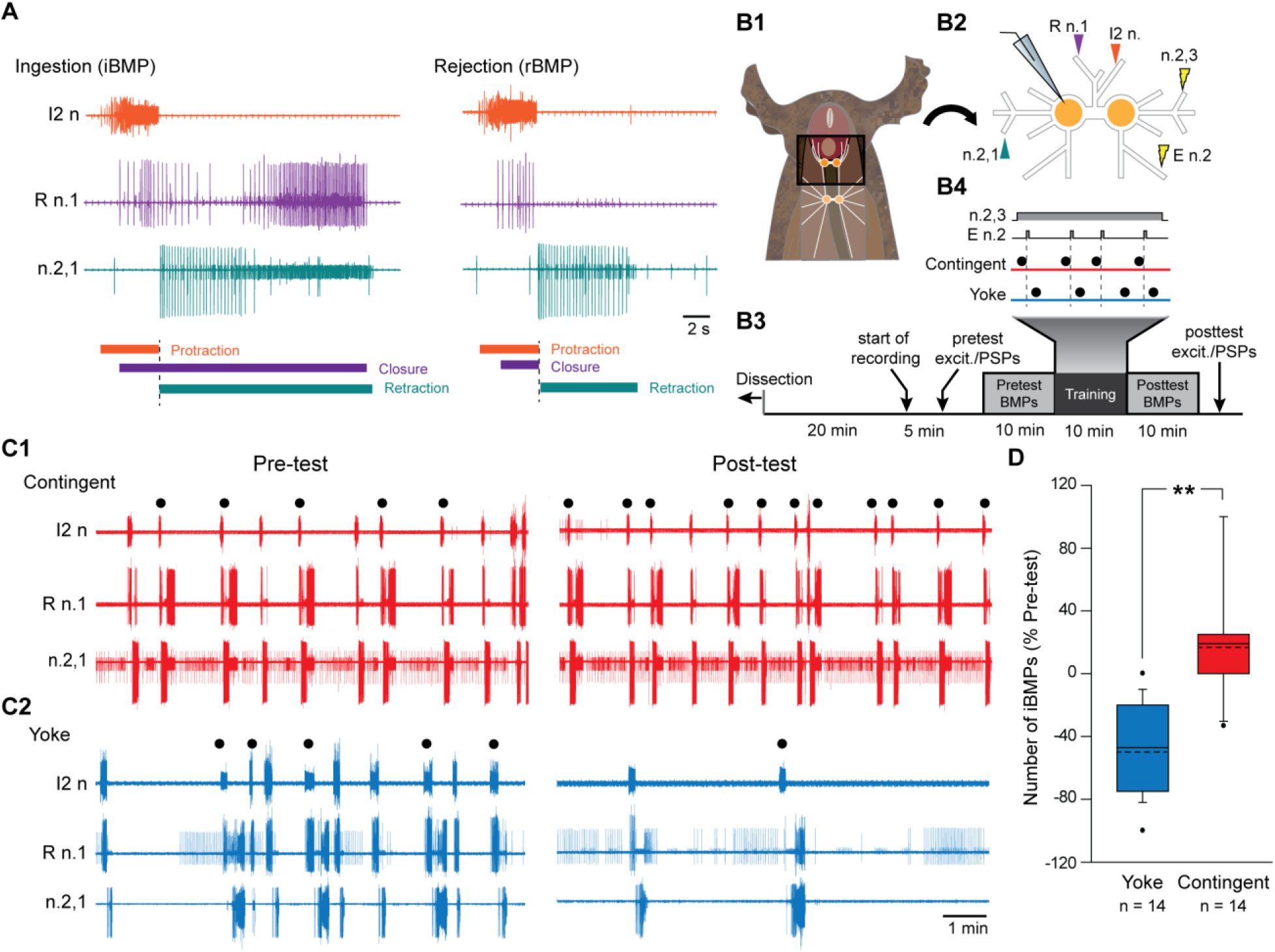
Buccal motor programs and their modulation by operant conditioning. The feeding CPG generates at least two types of patterned activity: one that mediates ingestion (iBMP) and one that mediates rejection (rBMP). ***A***, Recordings of BMPs produced by continuous, 2 Hz stimulation of Bn.2,3. The protraction phase was monitored via activity in I2 n. (orange trace and bar). The retraction phase was monitored by recording from Bn.2,1 (teal trace and bar). Closure motor activity was monitored via recording from R n.1 (purple traces and bar). ***A*** left, iBMP. The majority of R n.1 activity (> 50%) occurred during the retraction phase. ***A*** right, rBMP. Rn.1 activity was restricted in protraction phase. ***B***, Experimental design. ***B1***, Diagram of the animal with a longitudinal ventral incision. Nerves are white, ganglia are orange, and feeding musculature (i.e., buccal mass) is red. ***B*2**, *In vitro* preparation. Triangles indicate extracellular recording electrodes, lightning bolts indicate stimulating electrodes. ***B*3**, Timeline of the experiment. ***B4***, Training procedure. Closed circles indicate iBMPs, dotted line indicates start of rewarding En.2 stimulation. **C**, Recording of BMPs during 10 min pre- and post-test from a Contingent group (***C1***) and Yoke group (***C2***). In contingent group, the number of iBMPs was increased after the training, and decreased in yoke group. Filled circles indicate iBMPs. ***D***, Summary data. In this and subsequent illustrations: black bar in each box, median; dashed line, mean; box, interquartile range; whiskers, 10th and 90th percentile; filled circles, data points outside the 10th and 90th percentile; *, *p*<0.05; **: *p*<0.001.

**Figure 2.**
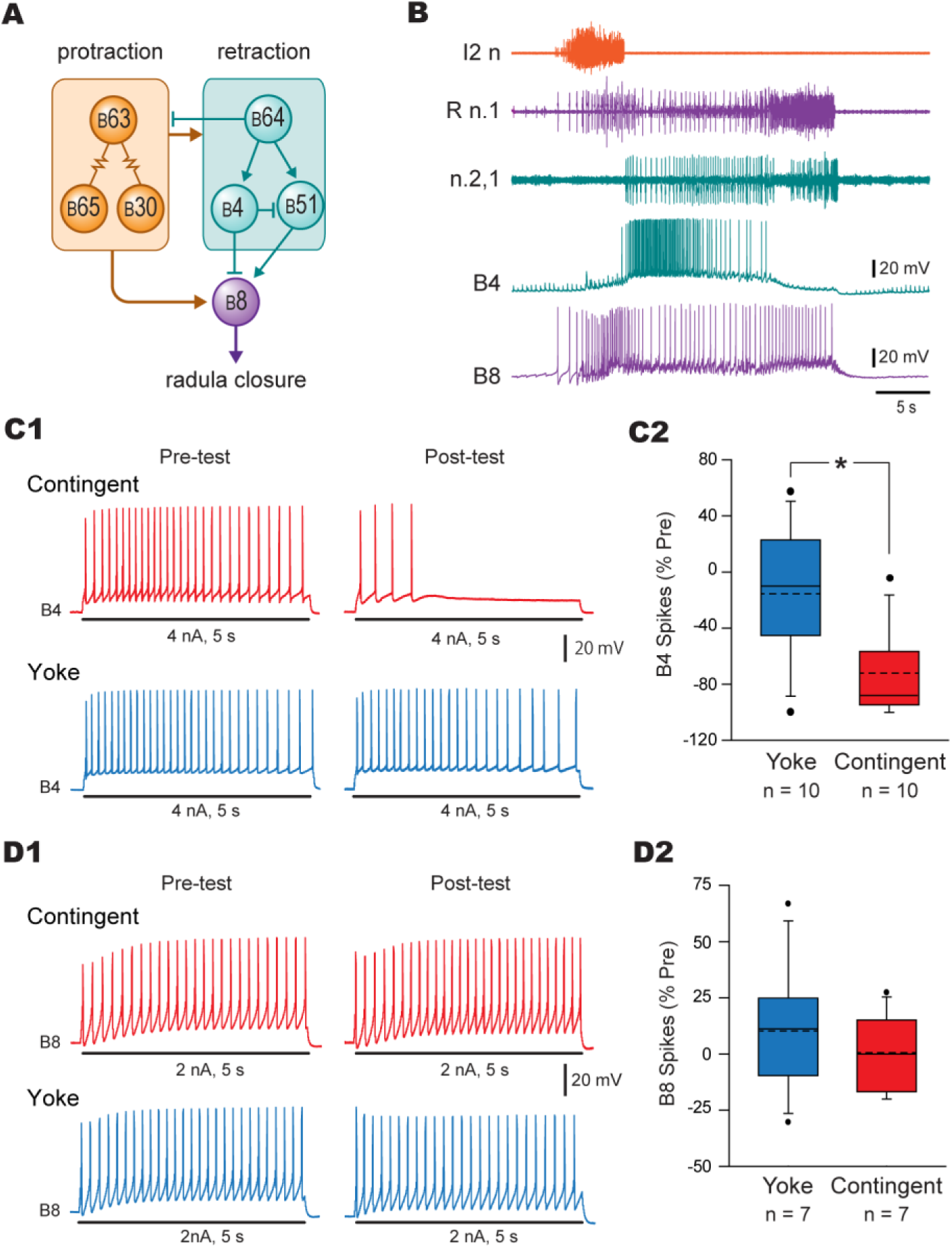
*In vitro* analog of operant conditioning reduced the excitability of B4 but not B8**. *A***, Simplified schematic of the feeding CPG. Activity in some neurons (e.g. B30, B63, B65) underlies protraction (orange), whereas activity in other neurons (e.g. B64) underlies retraction (blue). B51 plays a role in the expression of iBMPs through its excitatory connection to B8. Conversely, B4 inhibits B51 and B8. Thus, the level of activity in B4 plays a role in the expression of iBMPs. Arrows indicate excitation, whereas bars indicate inhibition. ***B***, A typical recording of B4 and B8 during an iBMP. B8 activity corresponded to R n.1 activity. ***C***, Contingent training decreased the excitability of B4. ***C1***, Representative intracellular recording of B4 excitability. The excitability was measured before (pre-test) and after (post-test) training. Preparations received either contingent training (red traces), or yoked training (blue traces). ***C2***, Summary data. ***D***, Conditioning had no effect on the excitability of B8. ***D1***, Representative intracellular recordings illustrate measurements of B8 excitability before (pre-test) and after (post-test) contingent training or yoked control. ***D2*.** Summary data.

### In vitro analog of operant conditioning

Animals were food-deprived for 1-3 d before the experiment and fed a piece of seaweed 30 min before dissection, at which time animals were anesthetized by isotonic MgCl_2_ with a volume equal to half the animal’s body weight (Brembs et al., 2004). The protocol for the *in vitro* analog of operant conditioning has been described previously (Nargeot et al., 1999a). Briefly, the buccal ganglia were isolated and transferred to the recording chamber containing artificial seawater with a high concentration of divalent cation (HiDi-ASW) composed of (in mM): 330 NaCl, 10 KCl, 30 CaCl_2_, 90 MgCl_2_, 20 MgSO_4_, 10 HEPES (pH was adjusted to 7.5 with NOH) (Nargeot et al., 1997) (Fig. 1*B1, B2*). HiDi ASW was used to reduce neuronal activity during dissection. The right hemi-ganglion was desheathed on the rostral side. Immediately after desheathing, the chamber solution was changed to normal ASW composed of (in mM): 450 NaCl, 10 KCl, 10 CaCl_2_, 30 MgCl_2_, 20 MgSO_4_, 10 HEPES (pH was adjusted to 7.5 with NOH) (Nargeot et al., 1997) and suction electrodes made with polyethylene tube for extracellular recording were used to monitor nerve activity. The preparations were at rest 20 min before the recording (Fig. 1*B3*) and maintained at 13-15°C by means of a Peltier cooling device for the duration of the experiment.

First, pre-test measurements of input resistance (R_in_), excitability, and synaptic strengths were made following a 5 min rest period after the beginning of intracellular recordings (see below). Then, sustained rhythmic BMP activity was elicited by continuous low-frequency stimulation of contralateral nerve n.2,3 (0.5 ms, 2 Hz) (Nargeot et al., 1997). Stimulation of En.2 (0.5 ms, 10 Hz), which contains dopaminergic afferents to the buccal ganglia, served as reinforcement during training (Fig. 1*B2* and 1*B4*) (Nargeot et al., 1997, 1999a,c; Kabotyanski et al., 1998; Martinez-Rubio et al., 2006). The intensity (2-4 V) of nerve stimulation of n.2,3 was adjusted so 10 consecutive stimuli elicited one-for-one spikes in B4. The intensity (3-8 V) of En.2 stimulation was adjusted so that: 1) one-for-one EPSPs were elicited in B4 by three consecutive stimuli at 1 Hz; and 2) 1 - 4 BMPs were generated with En.2 stimulation of 10 Hz for 6 s. Experiments consisted of a 10 min pre-test period, a 10 min training period, and a 10 min post-training period (Fig. 1*B3*). The pre-training period was initiated 5 min after the first occurrence of a BMP by tonic n.2,3 stimulation. During the post-training period, which immediately followed the training period, n.2,3 stimulation was paused briefly and post-test R_in_, excitability, and synaptic strength were measured. Each experiment included two groups: a contingent reinforcement group that received En.2 stimulation immediately following the expression of iBMPs and a yoke control group that received En.2 stimulation that was uncorrelated to pattern expression (Fig. 1*B4*). The experiments were done sequentially with a yoke experiment following each contingent experiment. The sequence of En.2 stimulation of each contingent preparation served as the template for the sequence of En.2 stimulation for its corresponding yoke preparation.

### Cell identification

Neurons were identified by their relative size, location, and physiological characteristics. For example, B51 was identified based on its small size, apposition to B15, and characteristic plateau potential (Plummer and Kirk, 1990). B8 was identified by its lateral position, large size, and one-for-one spikes in the radula nerve (Nargeot et al., 1997). B4 was identified by its medial position, large size, and inhibitory inputs to B8 (Rosen et al., 2000) and B51 (Plummer and Kirk, 1990).

### Testing B4, B8 and B51 properties in ganglia

Intracellular recordings were performed using conventional fine-tipped glass microelectrodes filled with 2 M potassium acetate (resistance 7-15 MΩ), with signals amplified by an Axoclamp-2B and digitized with a Digidata1322a (Molecular Devices, Sunnyvale, CA). Simultaneous recordings were made from the presynaptic (i.e., B4) and postsynaptic neurons (B8 or B51). B4 and B8 were recorded with dual electrodes whereas B51 was recorded with a single electrode. Experiments on the B4-to-B51 connection were performed in a different set of preparations than were those on the B4-to-B8 connection. R_in_ was measured in B4 and B8 by injecting a 5 s, 5 nA hyperpolarizing current pulse. The excitability was measured as the number of spikes elicited during injecting a 5-s duration depolarizing current pulse. B4 and B8 differ in their baseline excitability, so the intensity of depolarizing current was 4 nA for B4 and 2 nA for B8.

Measurements in which < 3 spikes were elicited for measurement of excitability during pre-test were excluded in the analysis for both the contingent and its yoke pair. Intrinsic properties were measured at resting membrane potential in the absence of n.2,3 stimulation. For measurements of IPSPs, B8 was held at -100 mV and B51 at -70 mV. A rest period of at least 4 min preceded the measurements of IPSPs before and after training. The effects of operant conditioning on excitability, R_in_, V_rest_ (resting membrane potential), and IPSP were assessed by the percent change in value between post- and pre-test ({(Post-Pre)/Pre}*100).

### Cell culture

Culturing procedures for B4 followed those described in Brembs et al. (2002) and Lorenzetti et al. (2008). Briefly, ganglia from adult *Aplysia* were treated with Dispase® II (10 units/ml) (neutral protease, grade II) (Roche, Indianapolis, IL) at 35°C for ∼3 h and then desheathed. Fine-tipped glass microelectrodes were used to remove individual B4 cells from the ganglia. Each cell was plated on poly-L-lysine coated petri dishes with culture medium containing 50% hemolymph, 50% isotonic L15 (Invitrogen, Carlsbad, CA). L15 was made of 350 mM NaCl, 25 mM MgSO4, 11.4 mM CaCl2, 29 mM MgCl2, 10 mM KCl, streptomycin sulfate (0.10 mg/mL), penicillin-G (0.06 mg/mL), dextrose (mg/mL) and 15mM HEPES. The pH of the culture medium was adjusted to 7.5. Cells were allowed to grow for 4-5 days. Prior to recording, the culture medium was exchanged for a solution containing 50% ASW and 50% isotonic L15.

### Single-cell analog for B4

The procedures for the single-cell analog were similar to those established previously (Brembs et al., 2002; Lorenzetti et al., 2008). Neuron B4 in culture had a higher R_in_ and lower firing threshold (R_in_ = 8.1 ± 0.6 MΩ; threshold = 0.6 ± 0.1 nA, N = 10) than cells in ganglia (R_in_ = 3.3 ± 0.1 MΩ; threshold = 3.0 ± 0.2 nA, N = 10), which is presumably due, at least in part, to the absence of the electrical coupling in the isolated neurons. R_in_ was measured by injecting a 5-s duration -1 nA hyperpolarizing current pulse. Excitability of cultured B4 was measured as the number of spikes elicited by a 5-s duration 1.5 nA depolarizing current pulses. The membrane potential was held at -70 mV, which was slightly more negative than the average resting potential (−63 ± 0.7 mV) of B4 in ganglia. Dopamine was iontophoresed through a fine-tipped glass microelectrode (resistance 10-15 MΩ) (Fig. 3*A*). The concentration of DA in the electrode was 200 mM with an equal concentration of ascorbic acid to reduce oxidation of DA. A retaining current of -3 nA in the DA electrode was used during the course of the experiment. The current was transiently stepped to +1 nA (6-s duration) to eject DA.

**Figure 3.**
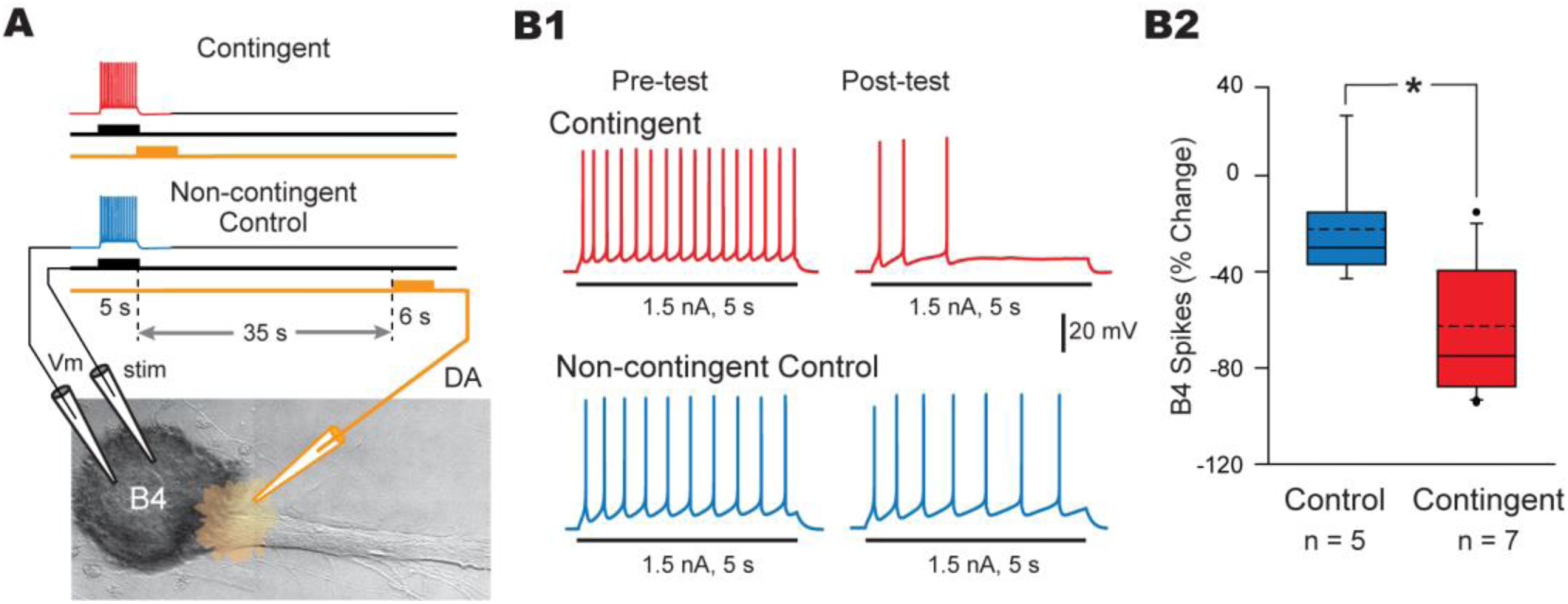
Single-cell analog of operant conditioning reduced B4 excitability**. *A***, Diagram illustrating the protocol for the single-cell analog of operant conditioning. In the contingent group, DA was applied extracellularly to the axon hillock region by iontophoresis immediately following a train of spikes in B4 that was elicited by a 5-s suprathreshold depolarizing current injection, whereas in the non-contingent group DA was applied 35 s after stimulation of B4. ***B***, Contingent DA application decreased the excitability of B4 in cell culture. ***B1***, Representative intracellular recordings illustrating measurement of excitability before (pre-test) and after (post-test) contingent DA application (red-traces) and non-contingent control (blue-traces). ***B2***, Summary data.

### Statistical Analysis

Statistical significance was set at P < 0.05 and all comparisons were two-tailed. Comparisons between two groups were made using paired t-tests or student’s t-tests for data that did not significantly differ from a normal distribution, which was tested by Shapiro-Wilk test. For non-parametric tests, samples were analyzed with Wilcoxon signed-rank tests. Statistical analyses were performed using Sigmaplot 12.0 (Systat Software, San Jose, CA).

### Computational model

This study used a conductance-based model representing a subset of CPG neurons sufficient to generate BMP-like activity elicited in isolated ganglia. The model is a modified version of that in Costa et al. (2020), implemented with the Simulator for Neural Networks and Action Potentials (SNNAP, version 8.1; Baxter and Byrne 2007). The methods used to develop the CPG model have been described previously (Ziv et al., 1994; Cataldo et al., 2006; Baxter and Byrne, 2007; Costa et al., 2020). Briefly, the CPG model includes sub-models of the cerebrobuccal interneuron CBI-2, six protraction neurons (B20, B31, B35, B63, B65, and B40), three retraction neurons (B4, B51, and B64), and a post-retraction neuron (B52). Neurons B31, B51, and B64 were modeled with two compartments (soma and axon), whereas the other neurons were modeled with a single compartment. The entire model has 49 excitatory, 32 inhibitory, and 22 electrical coupling synapses. Each synaptic connection was based on empirical data, with one exception. Because the mechanism of the transition between protraction and retraction is still under investigation, we included a fictive B63-to-B64 synapse, as used in Costa et al. (2020), to mediate the transition between these two phases. Conductance-based models of voltage-gated ion channels were added to each compartment, including only the necessary channels sufficient to simulate the unique firing properties of each neuron documented in the literature. Noise was a random variation in all conductances with an amplitude chosen to give a variety of patterns. 30 minutes were simulated, generating 56-118 BMPs depending on the parameter values of the OC plasticity loci. For experiments examining synergism 30-minute simulations were repeated 6 times. The simulations were concatenated and 100 BMPs were randomly sampled without replacement. This process was repeated 1000 times to obtain a distribution of the iBMP preference. The mean and standard deviation was calculated as an estimate of the width and central tendency of the distribution.

### Code Accessibility

The SNNAP files, equations, entire list of parameters and comparison to previous models is available at https://github.com/Byrne-Lab/momohara_neveu_2021 and http://modeldb.yale.edu/267112. The ModelDB access code is mn2021.

## Results

### Contingent reinforcement of iBMPs reduced the excitability of B4 but not B8

To evaluate the mechanisms of OC, this study used a previously developed *in vitro* protocol in which tonic stimulation of n.2,3 elicited a mix of ingestion-like and rejection-like buccal motor patterns (iBMPs and rBMPs) (Fig. 1*A*). As previously described (Nargeot et al., 1997), stimulation of En.2 was contingent upon the expression of iBMPs during training (Fig. 1*B4*). Figure 1*C* illustrates an example recording of BMP patterns expressed during pre- and post-test periods from two preparations, one in the contingent reinforcement group (Fig. 1*C1*), and one in the yoke control group (Fig. 1*C2*). Training led to a 17.8 ± 10.9 % (Mean ± SEM) ({(Post-Pre)/Pre}*100) change in iBMPs for preparations that received contingent reinforcement and a -47 ± 8.4 % change in iBMPs in a yoke group (Fig. 1*D*). There was a statistical difference between groups (Wilcoxon Signed Rank Test, W= 105, *P* < 0.001, n = 14 in each group). The number of pre-test iBMPs was not statistically different between yoke and contingent groups (contingent, 7.7 ± 0.8; yoke, 7.1 ± 0.9; paired t-test, t_(13)_ = 0.430, *P* = 0.674, n = 14 in each group). The decrease of iBMP frequency in the yoke group is consistent with observations in previous studies (Nargeot et al., 1997).

Previous studies found increased excitability of neurons B30, B63, B65, and B51 (Fig. 2*A*) (Nargeot et al., 1999a,b, 2009; Lorenzetti., et al., 2006, 2008; Mozzachiodi et al.,2008). To determine whether operant conditioning leads to a complementary decrease in excitability of a neuron that suppresses iBMPs, the intrinsic properties of B4 were measured (Fig. 2). A recording of B4 during an iBMP is shown in Figure 2*B*. Excitability was measured as the number of spikes during injection of an intracellular depolarizing current pulse (Methods). Contingent reinforcement led to a decrease in the excitability of B4 (−72.5 ± 10.9%) that was significantly different than the yoke control group (−15.5 ± 16.0%) (paired t-test, t_(9)_ = 3.330, *P* = 0.009, n = 10 in each group) (Fig. 2*C*). The excitability of pre-test was not statistically different between the groups (contingent, 25.0 ± 4.0; yoke, 22.5 ± 3.1; paired t-test, t_(9)_ = -0.517, *P* = 0.618, n = 10 in each group). The decrease in excitability was not due to changes in R_in_ (contingent, -27.1 ± 2.0%; yoke, -27.4 ± 2.1%; paired t-test, t_(7)_ = 0.090, *P* = 0.931, n = 8 in each group) or V_rest_ (contingent, 2.0 ± 4.0%; yoke, 2.9 ± 2.4%; paired t-test, t_(9)_ = -0.182, *P* = 0.859, n = 10 in each group).

We next investigated potential changes in the excitability of closure motor neuron B8 (Fig. 2*A*). B8 is a potential site of OC plasticity because an increase in its excitability could increase iBMP expression (Fig. 2*A*). In contrast to the changes in B4 excitability, no significant difference was observed in B8 excitability (Fig. 2*D*) between contingent (0.7 ± 6.8%) and yoke groups (10.2 ± 11.9%) (paired t-test, t_(6)_ = 0.871, *P* = 0.417, n = 7 in each group). There was no statistical difference in R_in_ (contingent, -2.0 ± 2.8%; yoke, -9.7 ± 6.8%; Wilcoxon signed rank test, W=5.0, *P* = 0.625, n = 5 each group) or V_rest_ (contingent, 2.7 ± 2.5%; yoke, 3.9 ± 3.3%; paired t-test, t_(6)_ 5.0, *P* = 0.584, n = 7 each group). These results indicate that operant conditioning reduces excitability of a specific subset of neurons (i.e., B4 but not B8).

### A single-cell analog of operant conditioning reduced B4 excitability

We hypothesized the OC-induced decrease in B4 excitability is intrinsic to B4 because it receives monosynaptic dopaminergic excitatory input from En.2 (Nargeot et al., 1999c), and iontophoretic DA responses in culture indicate dopamine receptors are expressed in B4 (data not shown). Therefore, we isolated B4 in culture and measured its properties before and after an analog of operant conditioning in which activity in B4 was contingent with application of DA, a known mediator of iBMP reinforcement (Nargeot et al., 1999c; Brembs et al., 2002; Lorenzetti et al., 2008). For single-cell analog preparations (Fig. 3*A*) two groups of B4 neurons were used, which differed in the delay between B4 stimulation and DA iontophoresis. The contingent group received DA iontophoresis (6-s duration) immediately following suprathreshold depolarizing current injections (5 s) into B4, whereas the non-contingent control group received DA 35 s after B4 stimulation. R_in_ and excitability were measured before and after conditioning (Fig. 3*B1*). The decrease in excitability of the contingent group (−62.7 ± 11.3%) was significantly greater than the non-contingent control group (−22.3 ± 12.1%) (t_(10)_ = 2.391 *P* = 0.038, n = 7 and 5 in contingent and non-contingent group, respectively) (Fig. 3*B2*). The pre-test excitability was not statistically different between the groups (contingent, 13.9 ± 2.1; yoke, 13.2 ± 3.0; paired t-test, t_(10)_ = -0.191, *P* = 0.852, n = 7 - 5 in each group). No significant differences were observed in R_in_ between the contingent (−22.1 ± 4.5%) and the non-contingent control group (−12.9 ± 3.7%) (t-test, t_(10)_ = -1.476, *P* = 0.171, n = 7 - 5 in each group).

These data indicate that the single-cell analog of OC reduced B4 excitability, and similar to *in vitro* conditioning, this decrease appeared to be independent of a change in input resistance. Moreover, these results indicate that the contingent-dependent decrease in B4 excitability was intrinsic to B4 and mediated by DA. We next examined whether the strength of chemical synaptic connections is altered by OC.

### An *in vitro* analog of operant conditioning reduced the B4-to-B51 IPSP, but not the B4 to B8 IPSP

B4 makes a large number of synaptic connections in the buccal ganglion (Gardner 1977; Gardner and Kandel, 1977; Fiore and Meunier, 1979, Susswein and Byrne, 1988; Kabotyanski et al., 1998; Sasaki et al., 2009; Dacks et al., 2012). Here, we focused on the B4-to-B51 and B4-to-B8 inhibitory synapses. These two connections could contribute to the changes in pattern generation associated with OC. A reduction in B4-to-B8 inhibitory synaptic connections would directly disinhibit B8, thereby favoring the expression of iBMPs. A reduction of the B4-to-B51 inhibitory connection could indirectly promote the expression of iBMPs by allowing increased activity in B51, and thus, increased activation of the B51-to-B8 excitatory connection (Fig. 4*A*).

**Figure 4.**
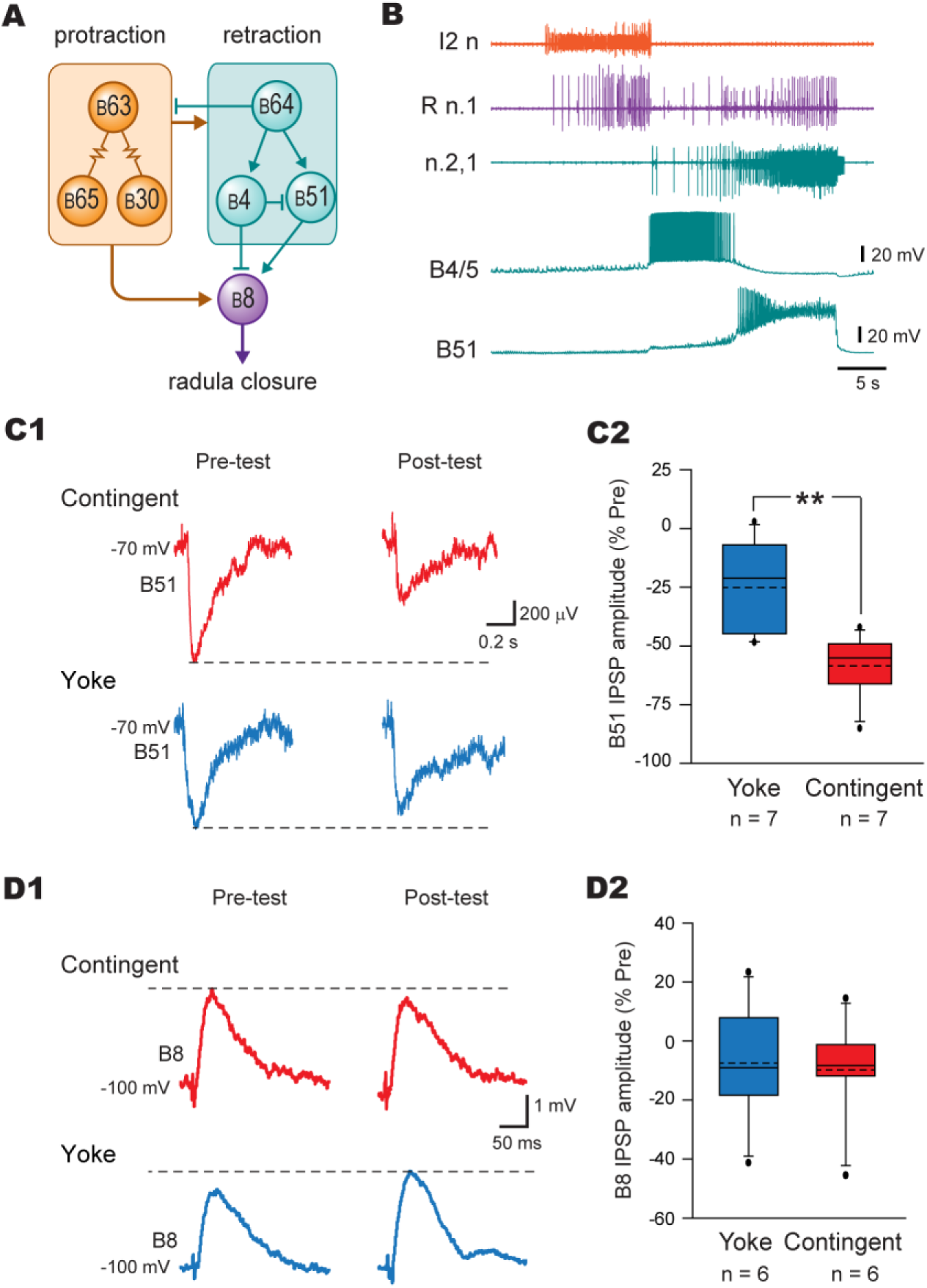
*In vitro* analog of operant conditioning reduced the B4-to-B51 synaptic connection but had no effect on the B4 to B8 synaptic connection**. *A***, Simplified schematic of the feeding CPG. ***B***, A typical recording of B4 and B51 during an iBMP. ***C***, Contingent training decreased the IPSP in B51. ***C1***, Representative intracellular recordings illustrating the measurement of the B4-to-B51 IPSP amplitude before (pre-test) and after (post-test) contingent (red-traces) and yoke control training (blue-traces). ***C2***, Summary data. ***D***, Conditioning had no effect on the IPSP in B8. ***D1***, Representative intracellular recordings illustrating the measurement of the B4-to-B8 IPSP amplitude before (pre-test) and after (post-test) contingent training (red-traces) and yoke control (blue-traces). Because the membrane potential was held at –100 mV, IPSPs were reversed in this panel. ***D2***, Summary data.

A simultaneous recording of B4 and B51 during a BMP is shown in Fig. 4*B*. The B4-to-B51 IPSPs from the contingent group were significantly reduced after conditioning as compared to the yoke group (Contingent, -58.4 ± 8.0%; Yoke, -25.1 ± 7.7%; paired t-test, t_(6)_ = 6.070, *P* < 0.001, n = 7 in each group) (Fig. 4*C*). The amplitude of the pretest IPSP was not statistically different between the two groups (contingent, 1.0 ± 0.1 mV; yoke, 1.1 ±0.1 mV; paired t-test, t_(6)_ = 0.834, *P* = 0.436, n = 7 in each group). These results indicate, for the first time, that OC of feeding behavior involves a change in chemical synaptic transmission as well as changes in intrinsic excitability and electrical coupling.

We next examined whether OC also affected the B4-to-B8 synapse (Fig. 4*A*). There was no difference in the B4-to-B8 IPSPs between the contingent and yoked group (Fig. 4*D*) (Contingent, -10 ± 8.0%; Yoke, -7.6 ± 9.0%; paired t-test, t_(5)_ = 0.154, *P* = 0.884, n = 6 in each group). These results indicated that plasticity is selectively induced among B4 chemical synaptic connections (e.g., plasticity at the B4-to-B51 connections, but not at the B4-to-B8 connection).

### A conductance-based model of the CPG

We hypothesized that a decrease in the excitability of B4 and in the B4-to-B51 inhibitory connection would contribute to biasing the CPG toward generating iBMPs. It is difficult, however, to estimate the relative contributions of B4 changes compared to previously discovered OC-induced loci of plasticity, such as increased excitability and coupling of neurons active during protraction that initiate BMPs (i.e., B30, B63, B65, B30↔B63, B63↔B65) (Nargeot et al., 2009; Sieling et al., 2014), and B51 that selects iBMPs (Nargeot et al., 1999a,b; Lorenzetti et al., 2006, 2008; Mozzachiodi et al., 2008). To address this issue, we used a conductance-based computational model of the CPG to examine the relative contributions of plasticity loci.

The CPG circuit model is illustrated in Fig. 5*A1*. Simulations of this model with all parameters set to control values generated activity resembling most of the features of physiological BMPs (compare Fig. 5*B* to Figs. 2*B* and Fig. 4*B*). Note the unique pattern of activation of each neuron during the simulated BMPs. The CPG also randomly switches between iBMPS and rBMPs (see Fig. 6). The variability was built into the model by introducing noise to the conductances. Consistent with literature, some neurons are more active during iBMPs (e.g., B8 and B51), whereas others are more active during rBMPs (e.g., B20, B34). Thus, this model reflects the selective engagement of subcircuits specific for each behavior. Taken together, these results suggested the model recapitulates the salient features of fictive feeding.

**Figure 5.**
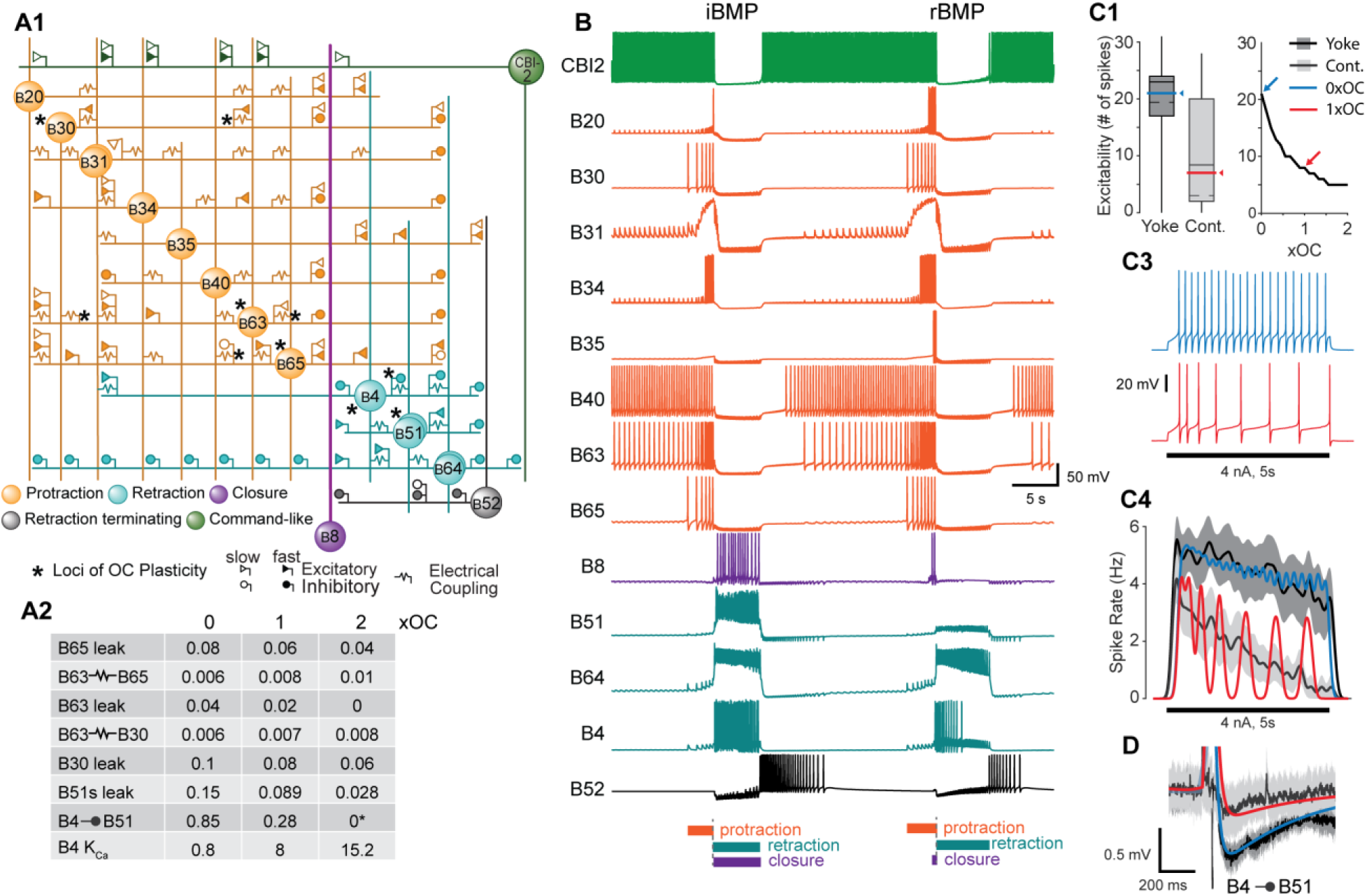
Overview of the CPG model and the OC plasticity loci. ***A1***, Diagram of the synaptic connections and the loci of OC plasticity (*) identified in this and previous studies. Neurons depicted with two circles have an axonal and soma compartment because these neurons exhibit plateau potentials. ***A2***, Table illustrating the values of parameters simulating OC-induced changes as compared to yoke-control parameter values. Column 0 represents the post-test yoke-control values, Column 1 represents the OC post-test-values. Column 2 OC represents hypothetical 2xOC post-test values mimicking learning. ***B***, Example of an iBMP (left) and rBMP (right) from a simulation where all parameters are set to yoke values. ***C1***, Excitability of B4 model for yoke control (blue) and contingent (red) conditions compared to empirical data for yoke (dark-grey) and contingent (light-grey) measured during posttest during a 5 s, 4 nA pulse (see *C3*). Color scheme the same for *C1-4*. ***C2***, Excitability for all parameter values used to modify B4 excitability. Test stimuli same as *C1* and *C2. **C3**,* Comparison of the time course of activity (Hz) during 4 nA current injection for empirical and model data. The model results are the mean of five simulation repetitions. Apparent oscillations emerged from the average of the five responses because of similar inter-spike intervals of the responses. Solid lines indicate arithmetic mean. Grey fill indicates SEM. ***C4***, Example simulation results showing the membrane potential trace during stimuli presented in *C3*. ***D***, Comparison of the IPSP waveform for empirical and model data.

**Figure 6.**
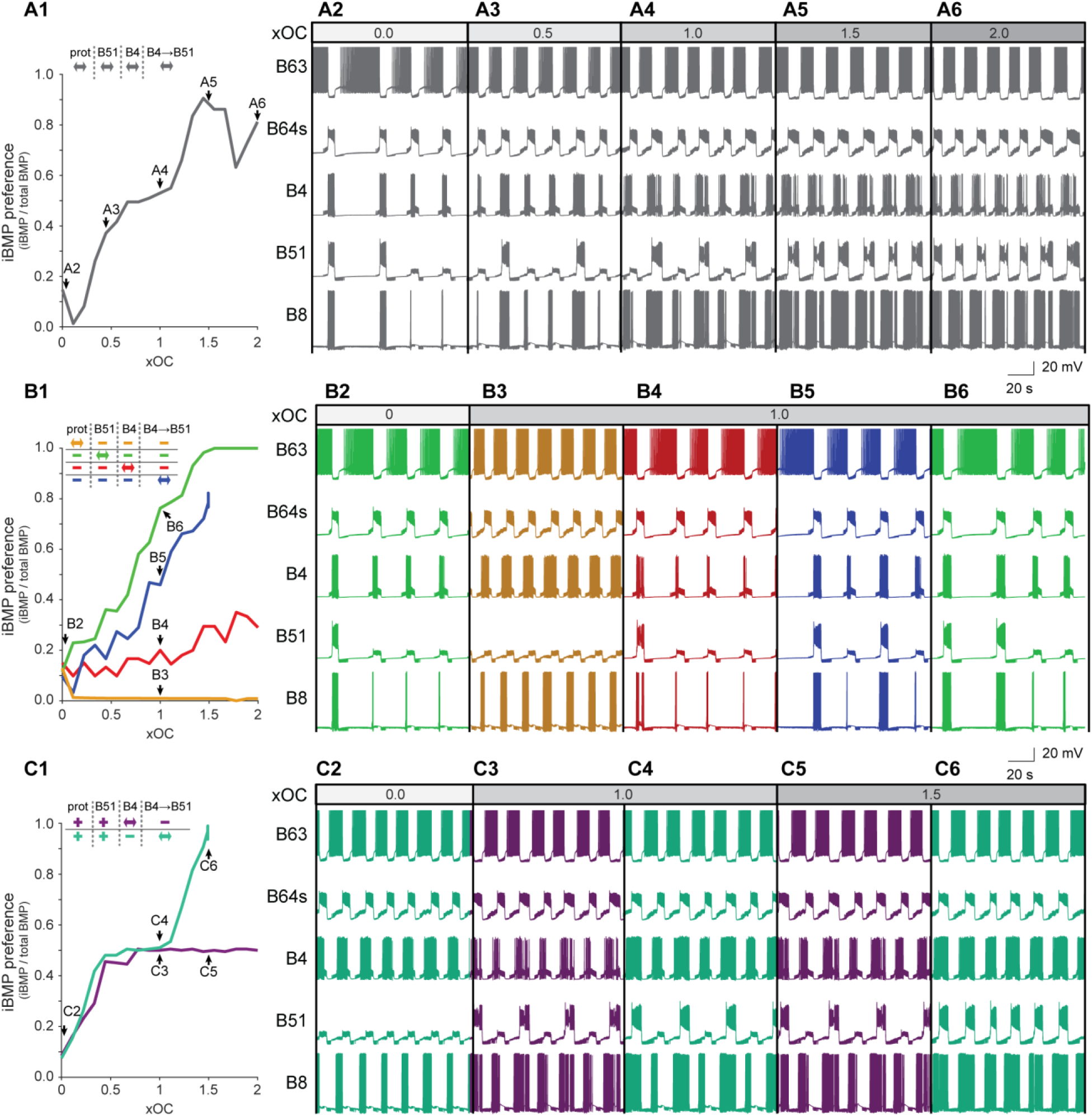
Plasticity loci have differing levels of efficacy to promote iBMP expression. ***A1***, Expression of iBMP preference for each independent simulation when the parameter values for all plasticity loci are set at varying levels relative to OC (see Fig. 5*A3*). Except where noted, 19 distinct parameter values were used to generate each line plotted in panels, *A1*, *B1* and *C1*. The letters on the lines correspond to the simulations in *A2-C6*. ***A 2-6***, Example traces from simulations in *A1*. ***B***, Expression of iBMP preference when only parameters for a single plasticity locus were modified at a time. For simplicity, all parameters modified by OC pertaining to protraction neurons (i.e., B30, B63, B65, B30↔B63, B63↔B65) were grouped into a single locus. The maximal conductance of the B4-to-B51 synapse was varied from 0.86 μS at Y to 0 μS at 1.5xOC and held at 0 μS for 4 additional simulations. ***C***, Expression of iBMP preference when protraction and B51 parameters were set to their 1xOC values and either the B4 or B4-to-B51 parameter was varied while the remaining parameter was held at yoke control value.

Contingent reinforcement increased iBMP expression compared to yoke training (Fig. 1). Therefore, we first tested whether concurrently modifying the parameters for all the above plasticity loci (Fig. 5*A2*) increased iBMP expression. To model OC-induced increases in excitability for B51, B30, B63, and B65, we decreased the leakage conductance of each cell. To model OC-induced increases in electrical coupling, we increased the coupling conductance of the B30↔B63 and B63↔B65 electrical synapses. For simplicity, parameter changes involving the protraction loci B30, B63, B65, B30↔B63, and B63↔B65 were always imposed simultaneously and not examined in isolation. To model OC-induced changes in B4, we added a Ca^2+^-activated potassium (K_Ca_) conductance to B4. The parameter for this conductance were taken from Baxter et al. (1999). This model reproduced the spike rate adaptation observed in B4 and increasing K_Ca_ maximum conductance above control values mimicked the decrease in excitability (Fig. 5*C*). The decrease in the B4-to-B51 IPSP was mimicked by decreasing the maximal conductance for this connection. We set the above parameters to values matching yoke control ganglia (Y values). OC was simulated by adjusting parameters to new values (C values) determined in this study and the literature. Parameter variations were obtained using the equation Y + A(C-Y), where A is a value between 0 and 2, and abbreviated as AxOC. Values of A above 1 examined whether further modification above posttest operant conditioning values enhanced iBMP expression. iBMP preference was calculated using the number of iBMPs divided by the total number of BMPs (iBMP/total BMP), which we denote as iBMP preference. When all parameters were set to yoke values (i.e., 0xOC) the simulated iBMP preference was 0.12 ± 0.01 (mean ± SEM, N = 4). Simulations of concurrent OC modifications at all loci increased the iBMP preference to 0.53 at 1xOC levels (Fig. 6*A4*). Further modification of all loci to 2xOC increased the preference to 0.81 (Fig. 6*A6*). These modifications also enhanced the rate of total BMPs, from 2.03 BMP/min at yoke parameter values to 3.33 BMP/min at 1xOC (data not shown). These simulations indicated that OC-induced plasticity increased total BMP expression and bias that activity toward the genesis of iBMPs. Moreover, these results indicated that more substantial modifications of these loci may further increase iBMP expression.

We next examined the ways in which alterations at individual plasticity loci change iBMP expression when the parameters for all other plasticity loci are set to yoke (i.e., 0xOC) values. Decreasing the strength of the B4-to-B51 connection alone increased the iBMP preference to 0.46 at 1xOC and to 0.78 at 1.5xOC (Fig. 6*B1, B5*). Note that at 1.5xOC the strength (g_max_) of B4-to-B51 was reduced to 0 μS. Decreasing B4 excitability alone led to a modest increase in iBMP preference to 0.20 at 1xOC and to 0.29 at 2xOC (Fig. 6*B1, B4*). Increasing B51 excitability alone led to the greatest increase in iBMP preference, to 0.76 at 1xOC and 1.00 at 2xOC (Fig. 6*B1, B6*). In contrast, concurrently modifying the parameters for the protraction loci towards their OC values led to a decrease in iBMP preference to 0.0094 at 1xOC and 0.0086 at 2xOC (Fig. 6*B1, B3*). Importantly, modifying the protraction loci enhanced the rate of total BMPs, from 2.07 BMP/min at yoke parameter values to 3.53 BMP/min at 1xOC (data not shown). These results indicate that increase of B51 intrinsic excitability, reduction of B4-to-B51 chemical synaptic strength, and to a lesser extent reduction of B4 intrinsic excitability can each increase iBMP expression, and that OC modifications of the protraction loci increase total pattern production while actually decreasing iBMP preference.

Given that changes of parameters at some loci towards their OC values increased iBMP preference, whereas changes at other loci decreased iBMP preference, we next examined what effect the plasticity loci of the current study (i.e., B4, B4-to-B51) had on iBMP expression when the parameters of the previously identified loci (i.e., increased excitability of B30, B51 B63, B65, and increased coupling of B30↔B63 and B63↔B65) are set to their 1xOC values. Setting the previously identified loci to their 1xOC values enhanced the effects of changes in the B4-to-B51 synaptic conductance and B4 excitability on iBMP preference (Fig. *6C*). For example, for B4 excitability, the enhancement of iBMP preference was only 0.20 at 1xOC when all other parameters were set to 0xOC (Fig. 6*B1, B4*), but 0.51 at 1xOC when the B51 and protraction loci were set to their 1xOC values (Fig. 6*C1, C3*).

### Synergism of the plasticity loci

The above results suggest potential synergistic effects between the different plasticity loci when modified combinatorically. To investigate this possibility further, we examined whether modifying a combination of parameters increased the iBMP preference to a greater extent than the arithmetic sum of iBMP preferences when parameters were modified individually. For reference we first set the protraction loci alone to 0.625xOC, which resulted in an iBMP preference of 0.010 ± 0.009 (Fig. 7*A, a*). We kept the protraction loci at 0.625xOC for all remaining simulations and then, individually or in combination, set the other loci to 0.625xOC, a value unlikely to have a basement or ceiling effect. When B51 excitability was modified the iBMP preference increased to 0.074 ± 0.024 (Fig. 7A*, b*). However, B4 excitability (Fig. 7*A, c*) or B4-to-B51 modifications (Fig. 7A*, d*) did not show a meaningful increase in iBMP preference in this circumstance (0.010 ± 0.009 and 0.012 ± 0.010, respectively), possibly because the decrease in iBMP preference caused by the protraction loci requires these sites to work synergistically to overcome the deficit. When B51 and B4 excitability were modified concurrently (Fig. 7*A, e*) the iBMP preference increased to 0.29 ± 0.04, which was much greater than the arithmetic sum of the iBMP preferences for these individual modifications (Fig. 7*A, b + c*) (0.074 + 0.010 = 0.084) This supra-additive effect on iBMP preference of combining the loci was also present when B51 was modified concurrently with B4-to-B51 change (Fig. 7*A, f*) (0.46 ± 0.04), compared to the sum of the individual loci (0.074 + 0.012 = 0.086). However, the supra-additive effect was marginal when B4 excitability and B4-to-B51 were modified concurrently (Fig. *7A, g*), resulting in an iBMP preference of 0.027 ± 0.015 compared to the sum of its constituents (0.010 + 0.012 = 0.022). We then compared the iBMP preference when all the loci were modified concurrently, with the effects of modifying only B51 and B4-to-B51 concurrently. Modifying all loci concurrently (Fig. *7A, h*) gave an iBMP preference of 0.46 ± 0.05 which did not exceed the summed iBMP preferences of B51 and B4-to-B51 modified concurrently (Fig. *7A, f*) and B4 excitability individually (Fig. *7A, c*) (0.46 + 0.01 = 0.47). This result indicates that B4 excitability has little added benefit on top of the B51 and B4-to-B51 synergism.

**Figure 7.**
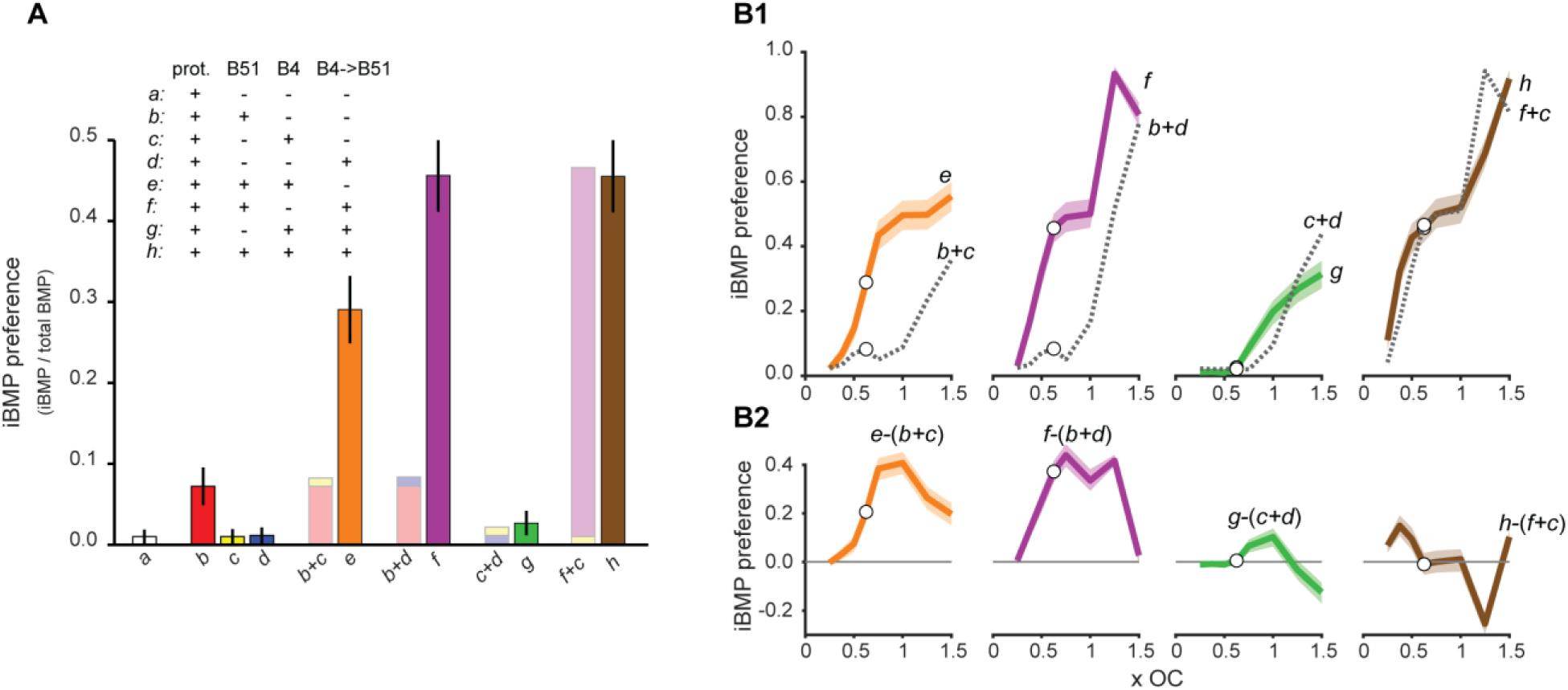
Synergism among sites of plasticity**. *A***, Example result of combinatorically adjusting the parameters at 0.625xOC values. Solid bars are results from simulations. Lighter bars with a combination of colors are arithmetic addition of the indicated colors. Mean and standard deviation of the mean are shown. Solid bars greater than the arithmetic sum of their parts indicate synergism between those loci. ***B1***, Results from simulations examining the range of parameter values that result in synergism when added in combination. Solid color lines are simulation results for the same combination as in panel *A*. Broken lines are arithmetic sums of the indicated simulations. Circles represent the example of the data in panel *A*. ***B2***, Subtractions of the results in panel *B1*.

Finally, rather than setting the parameters at 0.625xOC, we examined the range of values for which these loci acted synergistically. We set the parameters to values ranging from 0.25xOC to 1.5xOC using the same combinations mentioned above. For nearly all values tested, modifying the B4 and B51 excitability concurrently resulted in a greater iBMP preference than did the arithmetic sum of the individual modifications (Fig. 7*B*, *orange trace*). This effect was also seen when B51 and B4-to-B51 were modified concurrently (Fig. 7*B*, *violet trace*). However, modifying the B4 excitability and B4-to-B51 loci concurrently did not synergistically increase iBMP preference consistently compared to the sum of the individual modifications (Fig. 7*B*, *green trace*). In addition, modifying all of the parameters concurrently did not generate a greater iBMP preference compared to the sum of the iBMP preferences of B4 excitability increase and of concurrent B51 and B4-to-B51 modifications (Fig. 7*B*, *brown trace*). These results suggest that synergism only occurred in combinations including B51 excitability, and that this synergism was sufficient to produce the majority of the behavioral phenotype of OC.

In summary, these results indicated that the OC-induced decrease in the B4-to-B51 synaptic connection, and to a lesser extent the decrease in B4 intrinsic excitability, increased the pattern selection of the CPG towards iBMPs. Furthermore, when these loci were combined with previously discovered OC plasticity loci, they enhanced both the rate of BMPs and the iBMP preference. Finally, these plasticity loci are capable of working synergistically to enhance iBMPs. However, this synergism was loci specific in that not all combinations were able to synergistically enhance iBMP preference.

## Discussion

Although examples of neuronal correlates of OC have been found in many animal species (for reviews see Martin-Soelch et al., 2007; Mozzachiodi and Byrne, 2010; Cox and Witten, 2019; Nargeot and Puygrenier, 2019), the full extent of changes and how these changes interreact with each other has been challenging to investigate. The present study used *Aplysia* as a model system to extend understanding of OC by demonstrating: 1) OC decreased excitability of neuron B4; 2) OC decreased the strength of the chemical synaptic connection from B4 to B51; 3) OC-induced changes were site specific; 4) the relative contributions of these changes with a computational model; and 5) potential synergism between the plasticity loci.

### Reduced B4 excitability

Previous correlates of OC in *Aplysia* include increases in the excitability of neurons that initiate the behavior (Nargeot et al., 2009; Sieling et al., 2014; Costa et al., 2020), and increased excitability of a plateau generating neuron, B51, involved in selection of ingestion (Nargeot et al., 1999b; Brembs et al., 2002; Lorenzetti et al., 2008). Here, we found OC decreased the excitability of B4. This decrease appears to be intrinsic to B4 and not due, for example, to some change in a tonic modulatory circuit or in electrical coupling, because the excitability decrease can be mimicked by pairing B4 activity in an isolated neuron with application of DA (Fig. 3). Interestingly, pairing activity in B4 with DA leads to a decrease, whereas pairing activity in B51 with DA leads to increased excitability of B51(Nargeot et al., 1999b; Brembs et al., 2002; Lorenzetti et al., 2008) . This difference in responses indicates that common signals (i.e., spike activity and DA) presented contingently mediate opposite changes in B4 and B51. In mammalian systems, it is well known that DA modulation is mediated at least in part by differential expression of receptor subtypes, such as D1-like and D2-like receptors, which have different downstream mediators and lead to different cellular changes (Beaulieu and Gainetdinov, 2011; Liang et al., 2014) . The increase in B51 excitability is mediated by a signaling pathway similar to D1-like receptor activation (Lorenzetti et al., 2008). It is possible that the decrease of B4 excitability was mediated by another receptor subtype, such as a D2-like receptor. Alternatively, the same receptor pathway may ultimately modify different ion channel targets in B4 and B51.

### Decrease in the chemical synaptic connection from B4 to B51 and its specificity

Previously identified neuronal correlates of OC include increases in the excitability and electrical coupling (Nargeot et al., 1999a, b, 2009; Lorenzetti et al., 2006, 2008; Mozzachiodi et al., 2008; Sieling et al., 2014; Costa et al., 2020). The present study identified the first example of an OC-induced change in the strength of a chemical synaptic connection in the *Aplysia* feeding CPG. A reduction of the B4-to-B51 inhibitory connection would indirectly favor the expression of iBMPs by allowing increased activity in B51 (disinhibition), and thus, increased activation of the B51-to-B8 excitatory connection (Fig. 4*A*). Interestingly, the OC plasticity was branch specific, decreasing the B4-to-B51 synapse but not the B4-to-B8 synaptic connection. Branch specific homosynaptic depression was observed in the early studies of the feeding CPG by Gardner and Kandel (1977). Other examples of branch specific plasticity have been observed in *Aplysia* (Trudeau and Castellucci, 1993; Martin et al., 1997). Branch specific plasticity is ubiquitous. For example, in the hippocampus, a modulation of synaptic plasticity can be localized to specific dendrites in CA1 pyramidal neurons (for review see Edelmann et al., 2017).

### Relative contributions and synergism between the loci of plasticity

Synergism between plasticity loci has been suggested to be important for learning (Gao et al., 2012). The potential for synergism was examined in the current study by computer simulations of a conductance-based model derived from empirical biophysical data and cellular firing characteristics. Reduction of B4-to-B51 synaptic strength and to a lesser extent reduction of B4 excitability increased iBMP expression. More importantly, the B4-to-B51 and B4 excitability worked synergistically with a previously identified loci of OC, B51 excitability. Combining the decrease in B4-to-B51 efficacy with the increase in B51 excitability was more than five times more effective at producing the change in behavioral phenotype of OC. However, not all combinations acted synergistically, suggesting that some information pathways may be more suited to cooperate synergistically and mediate changes in behavior. In addition, synergism between plasticity loci may play an important role in the initial learning process, therefore future studies could examine how synergism facilitations OC learning. It will be important to examine synergism and its specificity among loci in other memory systems.

## Author contributions

Y. Momohara and C. Neveu made equal contributions to the work. Y.M. performed the in vitro electrophysiological studies; C.L.N performed the computer simulations; H-M.C. performed the B4 culture experiments; D.A.B contributed to the experimental design and analysis of the results; and J.H.B supervised all aspects of the project. All authors contributed to the writing of the paper.

## Acknowledgements

This work was supported by National Institutes of Health grant R01NS101356. The authors thank P. Smolen for detailed comments on earlier drafts of the manuscript and E Kartikaningrum for preparing cultures.

